# The post-translational modification profile of TAR DNA-Binding Protein (TDP-43) in platelets of patients with Alzheimer’s disease: An exploratory study for blood-based biomarker development

**DOI:** 10.1101/2023.01.29.526122

**Authors:** Qwynton Johnson, Mahan Hadjian, Alpha Bah, Sean Smith, Edina Kosa, Abdulbaki Agbas

**Affiliations:** College of Osteopathic Medicine, Department of Basic Sciences, Kansas City University, Kansas City, MO, USA; Heartland Center for Mitochondrial Medicine, Kansas City, MO, USA

**Author notes:** **Correspondence:** Abdulbaki Agbas.

**Keywords:** TDP-43, post-translational modification, platelets, Alzheimer’s disease, biomarkers

## Abstract

The assignment of blood-based biomarkers for neurodegenerative diseases is of great clinical value. Well-developed and validated blood-based biomarkers can serve in early diagnosis and prognosis as well as aid in patient screening when recruiting for clinical trials. We attempted to establish a portfolio for post-translationally modified TAR DNA/RNA-binding protein (TDP-43), a regulator of nuclear transcription, in platelet cytosol obtained from patients with Alzheimer’s disease (AD) comparing to age-matched healthy subjects and a disease control cohort. We aimed to identify the most prominent post-translational modifications of TDP-43 as an AD-relevant biomarker and to demonstrate that such an assessment can be performed in peripheral blood. We have isolated TDP-43 protein from human platelet cytosol utilizing an Immunoaffinity chromatography. The eluates were immunoprobed with a series of antibodies raised against post-translationally modified proteins. We employed a capillary electrophoretic immunoassay (CEI) to assess the phosphorylated TDP-43 profile. We observed that SUMOylation, phosphorylation, ubiquitination, and cysteine oxidation of TDP-43 are more prominent in platelet cytosol of AD patients as compared to control subjects. These studies will pave the way for identifying disease-specific TDP-43 derivatives that can be potential biomarkers for early diagnosis and the development of therapeutics.

## 1 Introduction

Alzheimer’s disease (AD) is the most common cause of dementia and is currently estimated to affect more than 5 million people in the United States, with an expected increase to 13.8 million by the year 2060. The typical clinical presentation is progressive loss of memory and cognitive function, ultimately leading to a loss of independence, inflicting a heavy personal toll on the patient and the family. The costs of care of patients with AD in 2010 were estimated at more than $172 billion in the United States alone. The annual cost is projected to inflate to one trillion dollars by 2050 unless disease-modifying treatments are developed (1). Attempts are being made to fill this demand, with the global market for AD treatment expected to grow from $4.8 billion in 2020 to an estimated $6.3 billion by 2025. That said, such attempts have been met with challenges; clinical trials of Alzheimer’s drugs ultimately failed more than 99% of the time between 2002 and 2012 (2). While treatment is multifaceted, a deterrent to developing disease-modifying therapies is the lack of biomarkers for early detection and evaluation. Therefore, well-validated and well-defined biomarkers of Alzheimer’s disease are needed. These biomarkers will improve the design of clinical trials, allow for development of more effective therapeutics, and offer the opportunity for prevention trials.

TAR DNA-binding protein (TDP-43) is a pivotal protein in the regulation of transcription and, by extension, cell homeostasis (3). There are several studies demonstrating that post-translational modifications of TDP-43 occur in a wide variety of neurodegenerative states in the central nervous system (CNS) where TDP-43 is associated with plaque formation (4-6). These plaques are insoluble by nature and can be histologically observed only in post-mortem tissues. Both overexpression and downregulation of TDP-43 is toxic for cells (7). Cells are unpredictable to process post-translationally modified TDP-43 and that of aggregated forms. This feature makes the therapeutic targeting of TDP-43 a challenging task. Identification of disease-specific post-translationally modified (PTM) TDP-43 profiles is necessary so that therapeutic intervention may be logically designed; hence, to establish chemically modified TDP-43 as a disease-specific and a validated blood-based biomarker is of importance. The PTM profile of TDP-43 may help to discriminate neurodegenerative diseases based on the progress stage. This will enhance our understanding of the pathological TDP-43 occurrence in neurodegenerative diseases.

We are working on developing a blood-based biomarker for neurodegenerative diseases such as AD, amyotrophic lateral sclerosis (ALS), and inclusion body myositis (IBM). Peripheral blood sampling provides easy access for obtaining repeated specimens using less invasive methods. Blood-derived platelets are a particularly attractive platform, providing an encapsulated environment for the cytosolic form of TDP-43 protein protected from the actions of several degradative enzymes such as proteases and phosphatases in blood serum/plasma (8). We have demonstrated the occurrence of hyperphosphorylation of TDP-43 in platelets (8-10) and others showed that plasma TDP-43 is largely or wholly restricted to platelets (11). Other potential PTMs with the capacity to produce toxic gain of function in the cell, such as cleavage, aggregation, acetylation, ubiquitination, SUMOylation, and cysteine-oxidation, need to be documented since neurodegenerative diseases vary in their repertoire of PTMs of target proteins (12). Blood-derived platelets provide a surrogate venue with a less invasive access to the central nervous system to evaluate PTMs of TDP-43 during the course diseases.

In this explorative study, we will compile a PTM profile of TDP-43 in platelets derived from patients with AD. This approach, if successful, can be applied to other neurodegenerative diseases such as amyotrophic lateral sclerosis Parkinson’s disease, Niemen-Pick C, and motor neuron disease.

## 2 Materials and Methods

### 2.1 Platelet cytosol preparation

Alzheimer’s disease patients were recruited through University of Kansas Alzheimer’s Disease Research Center (KUADRC). Blood samples were obtained from KUADRC under an approved IRB protocol (IRB#299644-3). All patients underwent annual Uniform Data Set clinical evaluations, which includes a battery of cognitive tests. The subjects included in this study were all males. The preparation of platelets was described in a previous manuscript (10). Briefly, approximately 8-9 mL of venous blood were collected over a vacutainer collection tube containing acid citrate dextrose (ACD) anti-coagulant reagent and brought to the laboratory within 1 hr. Platelet-rich plasma (PRP) was carefully separated from the buffy coat after centrifuging at 200 x g for 20 minutes. Prostaglandin I_2_ (PGI_2_) was added to PRP to prevent platelet aggregation (1 µL of 1mg /mL PGI_2_ per mL of PRP), gently mixed, and subjected to centrifugation (1,200 x g for 15 minutes) to pellet the platelets. The platelet pellet was washed with 1.0 mL of citrate wash buffer and subjected to an additional centrifugation (1,200 x g for 15 minutes) to eliminate possible red blood cell contamination. The pellet was then ruptured in 0.6 mL of rupture buffer and further subjected to sonication. The resulting platelet homogenate was centrifuged at 20,000 x g for 30 min at 4 °C to remove platelet membranes. The supernatant was aliquoted and stored at −80 °C freezer until use.

### 2.2 High Performance Immunoprecipitation (HPIP)

#### 2.2.1 Cross-linking anti-TDP-43 antibody to Protein-A-conjugated affinity media

Approximately 10 µg commercial anti-TDP-43 rabbit antibody (Proteintech Cat #10782-2-AP) was chemically cross-linked to Protein-A-conjugated affinity media pre-packed in a pipette tip (PhyTip Cat #PTR-92-20-01) (tip-affinity column). Cross-linking chemistry was performed in the presence of disuccinimidyl suberate (DSS), a non-cleavable cross-linking agent, according to an established method in our laboratory. The procedure of cross-linking the primary antibody to affinity media eliminates the co-elution of IgG antibodies with the target protein; hence, it increases the stability of antibody binding (13) and allows for reuse of the tip-affinity column up to two more times without losing antibody binding efficiency. We employed a fully automated multi-channel pipette equipped with software (PureSpeed, RAININ Cat #17013549) for the cross-linking process. A second tip-affinity column cross-linked with 10 µg home-prepared non-immune rabbit IgG antibody was used as a negative control.

#### 2.2.2 Immunoprecipitation of TDP-43

Platelet cytosols obtained from Alzheimer’s patients and age-matched controls were added to a deep-well plate and the automated multi-channel pipette was placed into position over the plate. Pre-programmed pipetting was initiated. The HPIP procedure included steps for equilibration, capture, wash, and elution. The TDP-43-enriched eluate was then neutralized with a small amount of 1M Tris-Buffer (pH 8.0). This eluate proceeded to SDS-PAGE or was stored at −20 °C for future use. Detailed information on the HPIP program utilized with the electronic pipette is provided in **Supplemental Table-1**.

### 2.3 Immunoblot Analysis

One hundred micrograms of platelet cytosol protein from the immunoprecipitation eluate (see step 2.2.2.) were loaded into a home-prepared one-well analytical SDS-PAGE gradient (4-20%) gel. Proteins were resolved in the electrophoretic field (100 V; 1hr) under reducing conditions and subsequently transferred onto PVDF membrane (116 mA; 1hr). Total protein staining was performed according to manufacturer instructions (Revert 700, Total Protein Staining, LI-COR Cat #926-11021) to assess the protein transfer efficiency and to normalize protein loading after completing the immunoblotting procedure. The membrane was destained using commercial reagent (Revert 700, Reversal Solution, LI-COR Cat #926-11013) and then blocked in a fish serum blocking buffer (Thermo Fisher, Cat #37527) for 1 hr at RT. The membrane was dissected into 10 equal strips and stored at 4 °C until use. Each strip was individually incubated with a relevant primary antibody (i.e., anti-SUMO1 and -SUMO 2/3, anti-ubiquitin, anti-cysteine, and anti-acetyl-lysine antibodies) overnight at 4 °C. The following day, the individual membrane strips were thoroughly washed with Tris-Buffered Saline, 0.1% Tween-20 (TBST) buffer and incubated with IR-conjugated secondary antibodies (IRDye 800CW, Cat #926-32211 [rabbit], #926-32210 [mouse]) for 1 hr at RT. The membrane strips were washed with TBST for 3×5 min and 1×5 min with TBS to eliminate the nonspecific fluorescence generated by Tween-20. The membranes were then scanned by an imager (LI-COR Odyssey) and the images were analyzed by Image Studio™ Lite Quantification software (Ver.3.1.4).

### 2.4 Capillary Electrophoretic Immunoassay (CEI)

Anti-phospho-(S409/410)-TDP-43 antibody (Proteintech Cat #80007-1-RR) was used to determine the phosphorylated TDP-43 profile in both AD and ALS platelet cytosols. The assay was performed according to a well-established methodology developed in our lab (10). The samples were prepared according to manufacturer instructions (Proteinsimple kit Cat #SM-W004). The fully automated CEI analyzer (Proteinsimple, Cat#004-600. This product is currently discontinued. Updated versions are available) is capable of measuring the phosphorylated TDP-43 levels in very small volume (0.4 µL), precisely analyzed the results, and created the final graph. Due to this features, CEI became a preferred method in our laboratory for high throughput screening for protein-based analysis in platelet cytosols obtained from patients with neurodegenerative diseases.

### 2.5 Sequence based prediction analysis

The TDP-43 amino acid sequence (UniProt/Swiss-Prot: Q13148) was explored via several proteomic databases for all predicted post-translational modifications that can occur (14). We showed only the predicted PTM sites in human TDP-43, rendered by PhosphositePlus (v6.6.0.4)

## 3. Results

We chemically cross-linked the polyclonal anti-TDP-43 rabbit antibody and non-immune rabbit IgG on Protein-A-conjugated affinity material (**Fig.1**) and checked for any IgG cleavage during the washing and elution process. We did not observe IgG molecules in the eluates (**Supplemental Fig-1A**). One hundred micrograms of platelet cytosolic protein mixture was applied to the HPIP tip-affinity column and antibody-bound proteins were eluted. The immunoblots of the eluates obtained from the platelet cytosol of AD patients and age-matched control subjects showed different levels of PTMs of TDP-43. The signal intensities were normalized to total protein staining according to the manufacturer protocol and the data were graphically illustrated (**Fig.2A**). Ubiquitinated and SUMOylated TDP-43 appeared to be the most notable PTMs, expressing the greatest signal intensity. We have observed reduced cysteine oxidation of TDP-43 in platelet cytosol samples obtained from patients with AD (**Fig.2B**) as compared to age-matched controls (**Fig.2C**). We were not able to detect acetyl-lysine modification. Due to low protein concentration (1-1.5 mg/mL) of individual platelet cytosol, we pooled both AD (n=10) and age-matched control cohort (n=10) samples and performed HPIP only once as a proof-of-concept. For this reason, statistical analysis was not performed at this time (**Fig.2A**). Data obtained from the CEI assay revealed two phosphorylated TDP-43 (pTDP-43) species observed in platelet cytosol samples obtained from AD patients and from that of ALS patients (the disease control). There was a notable elevation of the pTDP-43 band localized at approximately 60 kDa in the AD platelet cytosol (green peak) as opposed to the healthy control (blue peak) and the ALS disease control (red peak). Note that the same anti-phospho-(S409/410)-TDP-43 antibody recognized different phosphorylated TDP-43 protein species in different neurodegenerative diseases (**Fig.3**). Although we have utilized a number of publicly available programs, the most popular PTM studies were revealed using the PhosphositePlus (Ver.6.6.0.4) online prediction program with a number of references (14). Peak heights on y-axis refers to how many publications are available in particular PTMs of TDP-43 (**Fig.4**). The non-immune rabbit IgG tip-column, assigned as a negative control, did not pull down TDP-43 from the platelet cytosol samples (**Supplemental Fig-1C**).

**FIGURE-1:**
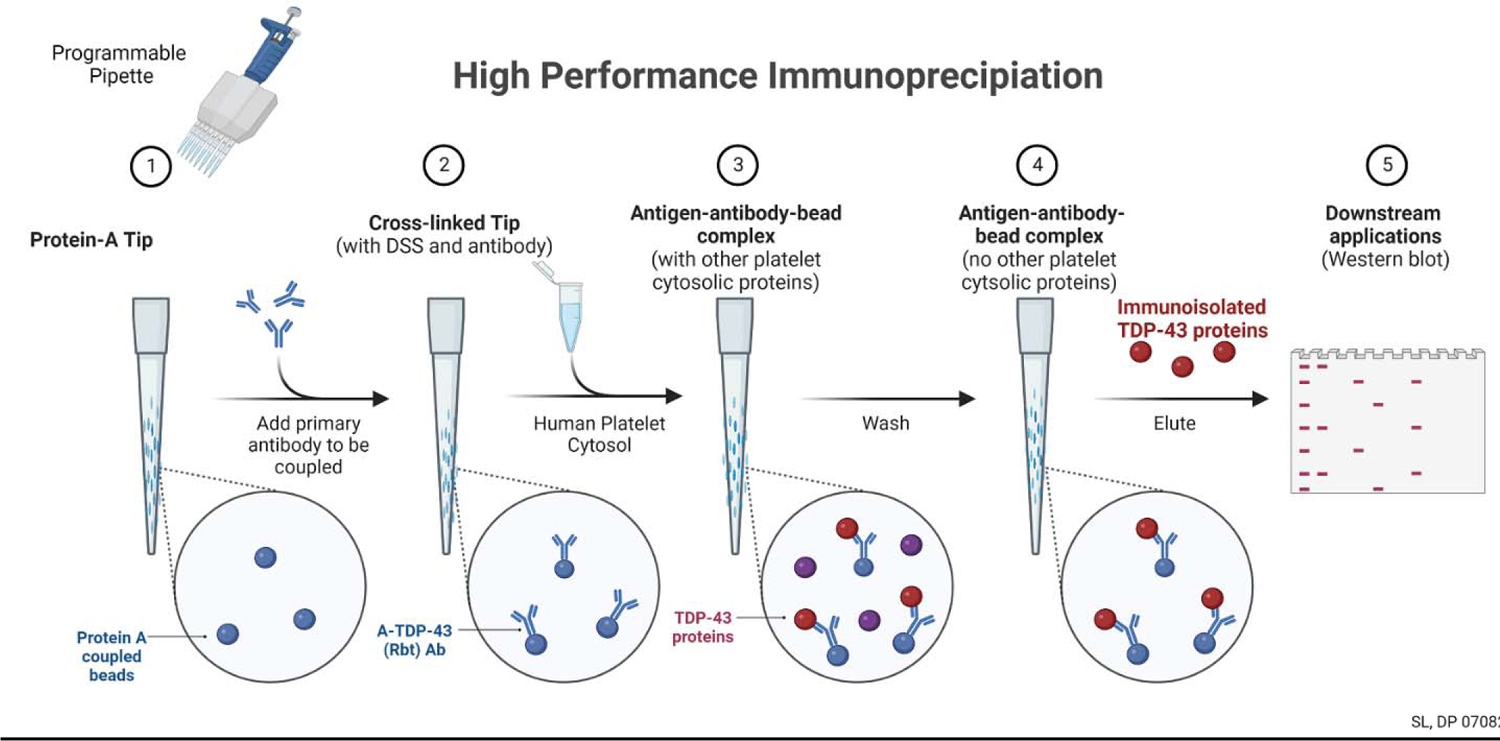
Illustration of the workflow for high performance immunoprecipitation (HPIP). Tip-columns were prepared by cross-linking a-TDP-43 antibody (steps-1,2) and used for high efficiency immuno-isolation of TDP-43 (steps-3,4)from platelet cytosol. The eluate were analyzed by classical immunoblotting assay (step-5). Created with BioRender.com

**FIGURE-2:**
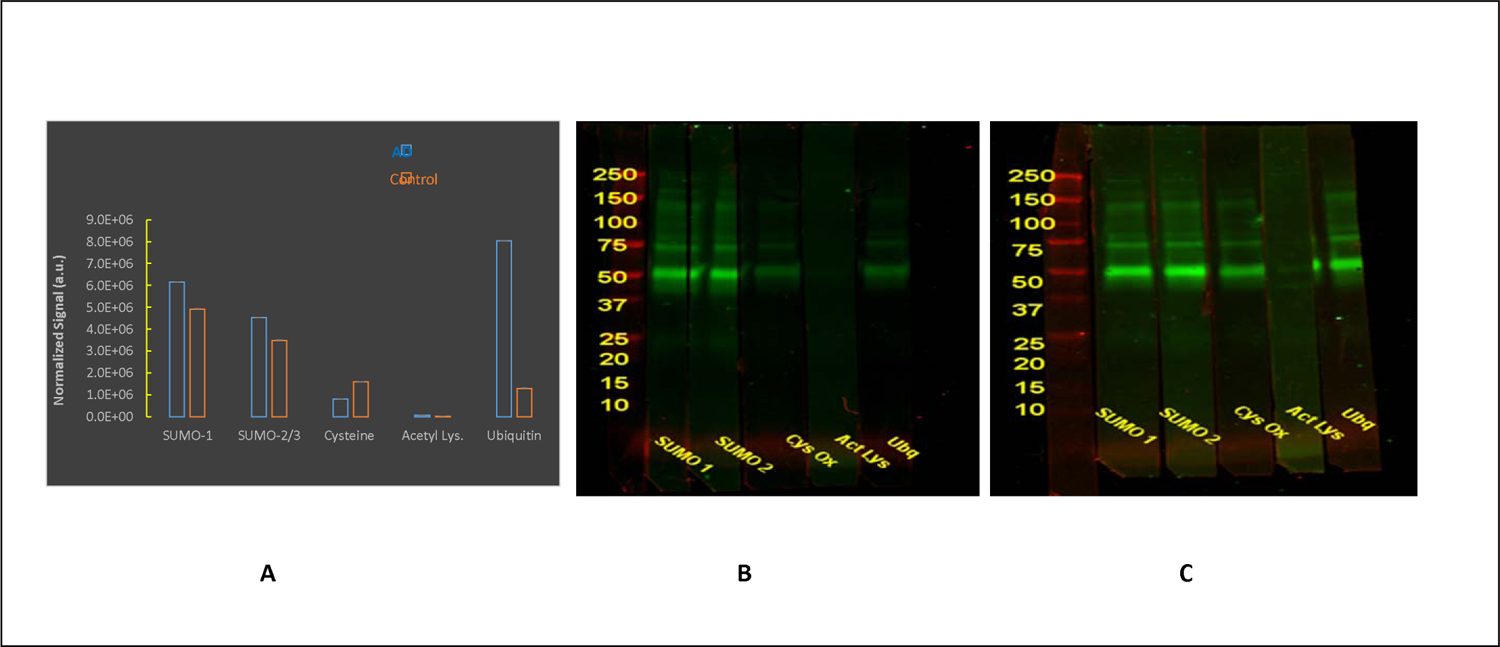
Post-translational modifications profile of TDP-43. **(A)** The bar graphs illustrate comparison between AD patients and age-matched control cohort in terms of PTMs of TDP-43 profile in platelet cytosol. The more noticeable elevations (blue bar) were observed in SUMOylation and ubiquitination detected by anti-SUMO-1/anti-SUMO2,3 and anti-ubiquitination antibodies, respectively. Since pooled platelet cytosol samples from AD patients (n=10) and age-matched control cohort (n=10) were used for HPIP, the assay performed only once; hence, no statistical analysis were performed. **(B)** Shows the immunoblot analysis of AD patients and **(C)** shows that of age-matched control cohort of PTM TDP-43 profile with various antibodies.

## 4. Discussion

The profile of PTMs of TDP-43 could be a useful indicator for the early diagnosis and prognosis of neurodegenerative diseases as well as discriminating such diseases from one another. Platelet-derived TDP-43 has been considered a potential candidate as a surrogate blood-based biomarker for neurodegenerative disease (8, 10, 15, 16). In this study, we analyzed the most popular PTMs of TDP-43 in platelet cytosols obtained from patients with AD as well as a healthy age-matched control and ALs as disease control group.

We chose platelet cytosol as the venue for measuring the TDP-43 profile. Platelets are anucleated, cell fragments derived from megakaryocytes (17-19). TDP-43 is a nuclear protein that has the ability to shuttle back and forth between the nucleus and cytosol (20). When TDP-43 proteins undergo PTMs due to disease, as in neurodegeneration, or physical trauma, such as repeated concussions in contact sports (21, 22), TDP-43 translocates to the cytoplasm. The cytoplasmic accumulation of TDP-43 is sufficient to induce aggregation through protein-protein interaction and phosphorylation (23, 24). Although remains largely unexplored, new studies support the notion that phosphorylation of TDP-43 occurs after aggregate formation in cytosol (reviewed in (25). Therefore, platelets are an appropriate system of study, reflecting a cytosolic environment where PTMs of TDP-43 will be abundant. In a separate pilot study, it has been shown that approximately 90 % of plasma TDP-43 was found in platelet enriched plasma as compared to 10% TDP-43 in platelet poor plasma (11) (R. Bowser, personal communication, October 12, 2022).

There are several publicly available PTM prediction programs that can be employed for forecasting possible PTMs of TDP-43. We have checked the majority of the sites provided in a review paper (14); that said, many of links were either inactive or under maintenance at the time of writing this manuscript. Of all the proteomic databases and tools that we tried, only phosphosite.org (https://www.phosphosite.org/homeAction.action) generated a comprehensive output. Therefore, we presented the data output of PhosphositePlus v6.6.0.4, which provided all popular PTMs along with references for each (**Fig.4**). Prediction analysis of TDP-43 for natural disorder regions (PONDR^®^) revealed that most of the phosphorylation events occur between amino acids 359-410 (**Supplemental Figure-2**).

We have isolated platelets from whole blood within a one hour timeframe by employing a two-step centrifugation as described in a previous manuscript (10). After lysis and removal of membranous fragments by centrifugation, the platelet cytosol samples were stored at −80 °C until use. We immunoprecipitated TDP-43 proteins from platelet cytosol obtained from AD subjects, as well as age-matched but otherwise healthy control subjects, employing HPIP technology as illustrated in **Fig.1**. Platelet cytosol samples obtained from patients with ALS were included as a disease control cohort for the CEI assay so as to compare the pTDP-43 profile in both diseases (i.e., AD and ALS). Traditional immunoblotting was employed to demonstrate that TDP-43 undergoes posttranslational modification in AD as compared to control subjects. One notable pattern observed in AD patients was a modest increase in TDP-43 SUMOylation as compared to healthy patients (**Fig.2**). The regulatory role of SUMOylation has been implicated in fundamental processes, with several reports indicating that the aging process could be influenced by SUMOylation (26-30). Based on our exploratory results (**Fig.2**), we may propose that the SUMOylation characteristics of TDP-43 could be considered as a disease-specific hallmark and possible target in AD. We recognize that we must also study other neurodegenerative diseases such as ALS, Parkinson’s disease, Huntington’s disease, multiple system atrophy, and progressive supranuclear palsy and demonstrate the SUMOylation pattern (or lack thereof) of TDP-43 in such diseases (31-33).

We have noticed that there is a pattern of increase in TDP-43 ubiquitination in parallel to increasing SUMOylation. These two PTMs vary in terms of function. Ubiquitination is used to tag proteins for degradation in proteasome machinery, whereas SUMOylation alters gene expression, chromatin structure, signal transduction, and maintenance of the genome (34). These two patterns (i.e., SUMOylation and ubiquitination) may suggest that TDP-43 ubiquitination is a response to aberrant protein expression caused by increased TDP-43 SUMOylation, an attempt to remove the aberrant protein via the proteasome apparatus. More work needs to be done concerning the SUMOylation characteristics of TDP-43 during AD progression into the more advanced stage. The other relevant biological question would be whether SUMOylation of TDP-43 induces proteasome activity so that ubiquitinated TDP-43 can be eliminated from cell.

The most obvious post-translationally modified TDP-43 species is pTDP-43, with 64 potential phosphorylation sites (41 Ser, 15 Thr, and 8 Tyr residue) available on the protein (**Supplemental Fig. 2**) (8). <u>**H**</u>owever, <u>**d**</u>isorder <u>**e**</u>nhanced <u>**p**</u>hosphorylation <u>**p**</u>redictor (DEPP) revealed that 20 Ser, 4 Thr, and 4 Tyr residues are statistically more probable sites for phosphorylation. Most of these sites are located in the C-terminus region of the sequence. We employed CEI technology, and observed that commercial anti-phospho-(S409/410)-TDP-43 antibody recognized two separate protein populations, at MW 47 kDa and MW 60 kDa, in platelets obtained from patients with ALS and patients with AD, respectively (**Fig.3**). This pattern has held for several ongoing clinical trials that we are involved in. It is currently unknown whether TDP-43 undergoes a hyper-phosphorylation process that prompts this presence in AD platelets. If that is the case, we may consider the ∼60 kDa pTDP-43 species to be an AD-specific signature protein discriminating from ALS, which instead expresses an alternate but equally specific pTDP-43 species at ∼47 kDa. More work needs to be performed in terms of reproducibility, specificity, and sensitivity of pTDP-43 measurements.

**FIGURE-3:**
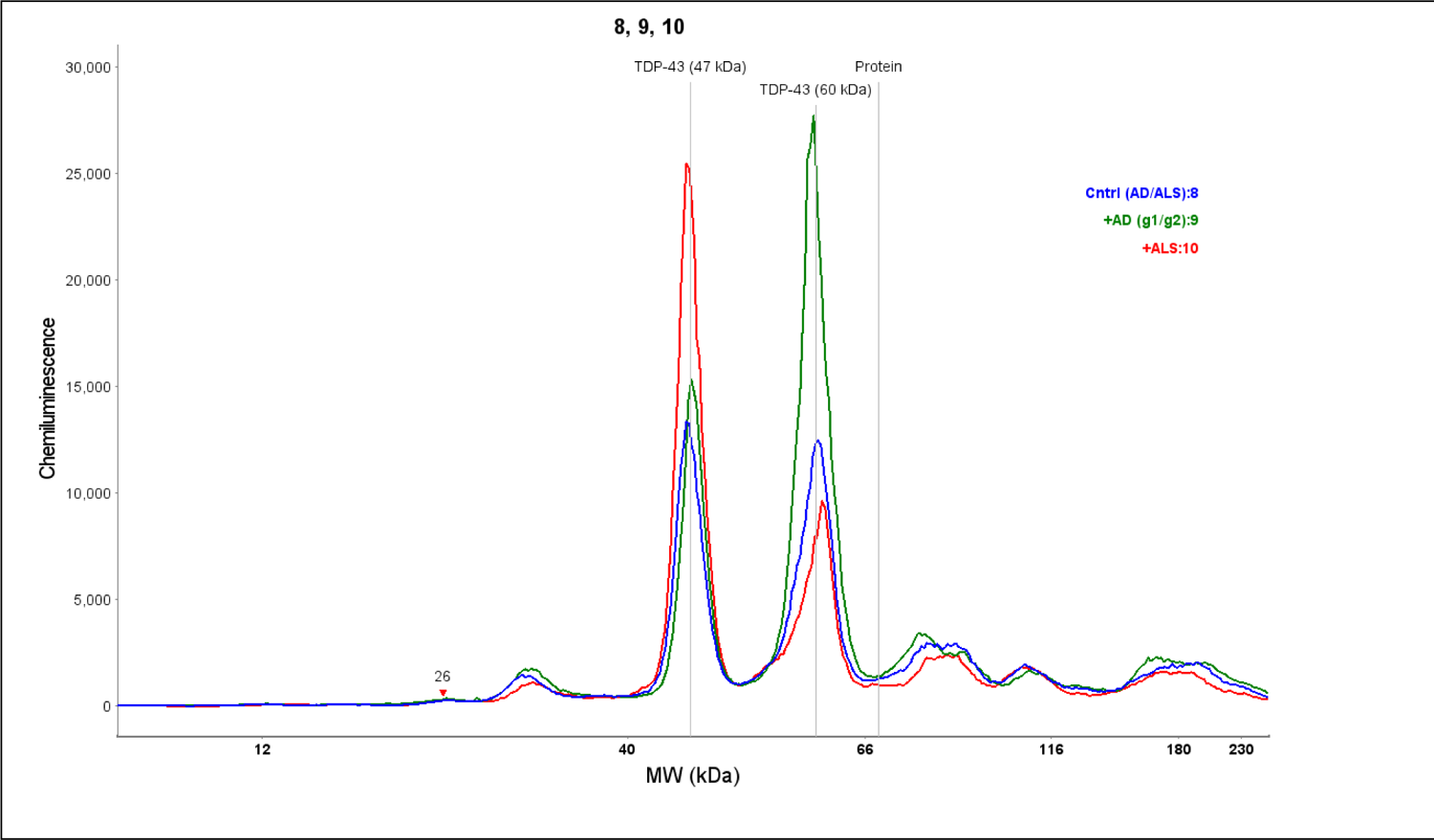
pTDP-43 electropherogram of capillary electrophoretic immunoassay (CEI). Anti-p(S409/410)TDP-43 antibody was used in this assay. The same antibody recognized TDP-43 species with different molecular weight. The prominent pTDP-43 peak appeared at ∼60 kDa (green color peak) for AD patients whereas the most prominent pTDP-43 peak appeared at ∼47 kDa (red color peak) for ALS patients. Control group pTDP-43 peaks (blue color peak) did appeared in both location with the same intensity.

**FIGURE-4:**
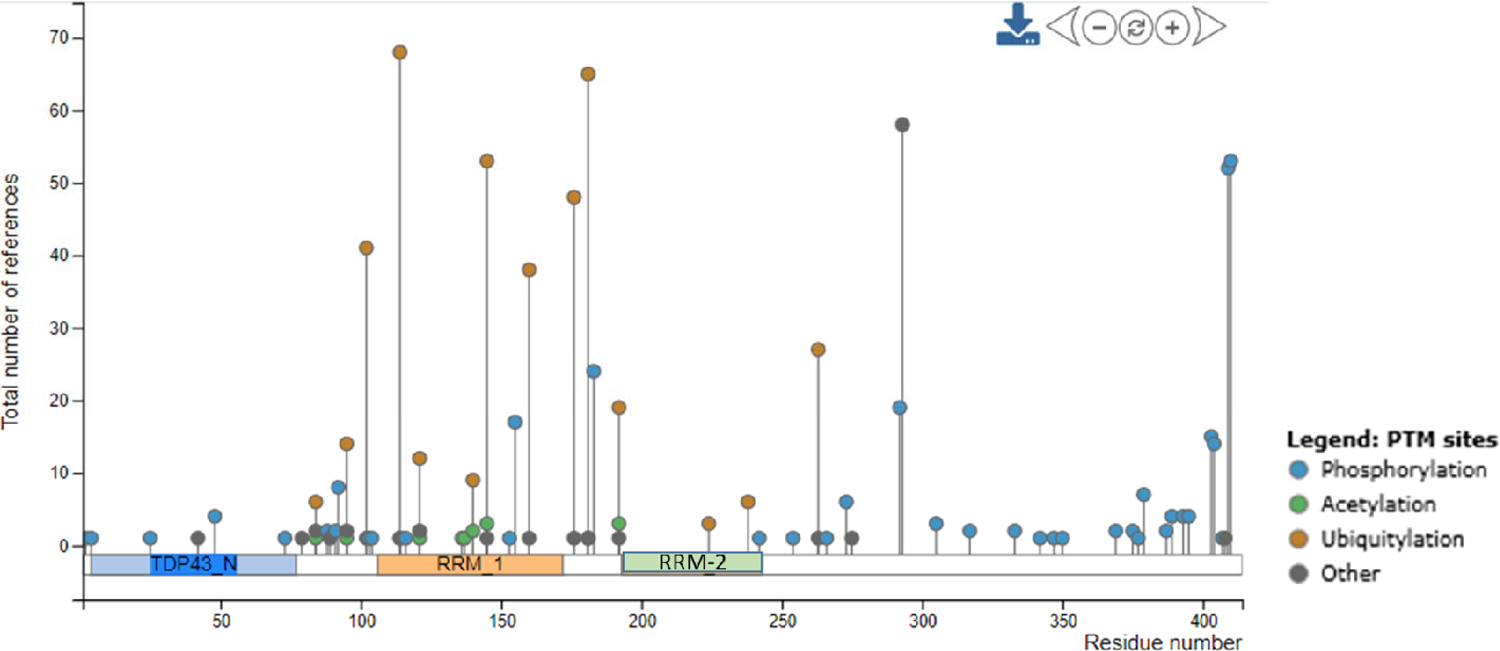
PhosphositePlus predictions fort post-translational modifications of TDP-43. TDP-43 amino acid sequence (Uniport/Swiss-Prot: Q13148) was uploaded in PhosphositePlus (Ver.6.6.0.4) program and several PTMs were predicted. The most notable PTMS were phosphorylation, acetylation, and ubiquitination. The height of circles represent the total number of published papers. The most phosphorylation sites were appeared to be in C-terminal region of TDP-43. The most ubiquitination sites were located on of the RNA-binding site of TDP-43 (RRM-1)

We have also observed that anti-phospho-(S409/410)-TDP-43 antibody recognizes cross-linked pTDP-43 in ALS at MW ∼230 kDa. (**Supplemental Figure-3**). It is possible that as a super-phosphorylated TDP-43 conglomerate, this TDP-43 species do not completely enter the separation matrix packed in the capillary column of CEI. High-stringency solubilization and reducing conditions may resolve some of the cross-linked pTDP-43 conglomerates, and we are currently working on this issue and testing whether AD platelet cytosol shows a similar pattern (i.e., ∼230 kDa protein conglomerate).

Analysis using CEI has precisely provided accurate molecular weight information for target proteins in every assay we have performed using commercial molecular weight markers. We did not use the other antibodies utilized during immunoblotting studies (i.e., anti-SUMO1, anti-SUMO2-3, anti-cysteine sulfonation, anti-acetylated Lys, anti-ubiquitin antibodies) in the CEI assay system. Not every antibody works well in the CEI platform and optimization steps (concerning kinetics of binding and minimizing background) must be taken for each antibody prior to use in CEI assays. We routinely measure pTDP-43 levels in human platelet cytosol for several clinical trials (16); hence the anti-phospho-(S409/410)-TDP-43 antibody was well optimized for CEI assays. We have included CEI data for only the pTDP-43 profile in this manuscript CEI studies utilizing other commercial antibodies may be done once the binding kinetics of those antibodies have been optimized in the future.

Acetylation of proteins can occur as a co-translational or post-translational modification (35). Two main sites (Lys145 and Lys192) of acetylation in TDP-43 were identified by generating acetylation mimics (36). However, in a contradictory study, mass spectrometry analysis revealed that residues Lys145 and Lys192 were not acetylated in TDP-43 inclusions found in brain samples of ALS patients (37). We were also unable to assess acetylation of TDP-43 on Lys residues by HPIP/immunoblotting methods (**Fig.2**). Either the anti-acetyl lysine antibody that we used for this study (Cytoskeleton, Inc., Cat# AAC01-S) did not work in our hands or there was no recognizable acetylation of TDP-43 in AD platelet cytosol. In the future, we will use antibodies from other vendors, paying special attention to other potential acetylation sites of TDP-43, such as Lys82 (37), to clarify this ambiguity regarding whether Lys acetylation in TDP-43 is disease-specific.

The other peculiar trend we noticed was that all forms of TDP-43 expressing oxidized cysteine moieties with an oxidized thiol group (i.e., sulfenic RSOH, sulfinic RSO_2_H, and sulfonic RSO_3_H) were reduced in AD patients as compared to age-matched controls. It seems that there was less cysteine modification in AD than in age-matched controls (**Fig.2**), which is contrary to what we expected (i.e., more sulfonated cysteine residues in TDP-43). It has been hypothesized that reactive oxygen species may trigger some TDP-43 pathology (38). Cohen et al., proposed that there are at least six cysteine residues (Cyst39, Cys50, Cys173, Cys175, Cys198, and Cys244) in the TDP-43 protein sequence that are targets for cysteine oxidation (38). We have demonstrated in a previous manuscript that functional protein undergoes aggregation via -SH groups in the aging brain (39).

Since a disulfide bridge (S-S) forms between two cysteine amino acids, we were expecting that cysteine modification in TDP-43 would be elevated. It is possible that TDP-43 takes on a more soluble form in platelet cytosol; hence, TDP-43 aggregation is less. This conjecture needs to be experimentally tested by first pre-treating isolated platelets with dimedone (5.5-Dimethyl-1, 3-cyclohexanedione), which will help identify cysteine residues expressing sulfenic acid groups from other cysteine oxoforms (40), then immunoprecipitating TDP-43, and immunoprobing with an anti-cysteine (sulfonate) antibody.

In conclusion, several post-translational modifications of TDP-43 can be analyzed in human platelet cytosol, mediated by the compromise of the blood-brain barrier in neurodegenerative diseases (41, 42). Many CNS biomolecules can be observed in peripheral blood tissue; therefore, the profile of platelet TDP-43 and its derivative species could serve as a surrogate biomarker for neurodegenerative diseases.

## 3 Conflict of Interest

The authors declare that the research was conducted in the absence of any commercial or financial relationships that could be construed as a potential conflict of interest.

## 4 Author Contributions

QJ, MH, AB, and SS are medical students. All were involved in experimental procedure, data analysis, and graphic creations. EK helped the students perform the experiments. MH was involved in editing the manuscript. AA conceived the idea, planned the experiments, and wrote the manuscript.

## 5 Funding

This project is funded by an intramural grant awarded to AA.

## 6 Acknowledgments

The authors acknowledge the Office of Research and Sponsored Programs at Kansas City University for awarding summer research fellowships for QJ, MH, and AB.

## FIGURE LEGENDS

**Supplemental Figure-1: Anti-TDP-43 antibody cross-linking onto tip-affinity column. (A)** After cross-linking anti-TDP-43 antibody (rabbit), a HPIP procedure was performed with PBS buffer (pH 7.4) as input sample, Flow-through and eluate was collected and analyzed by 4-20% gradient SDS/PAGE. The gel was stained by silver staining method. No IgG leaching was observed, meaning we achieved to cross-link the antibody to tip-affinity column. **(B)** Anti-TDP-43 antibody was chemically cross-linked onto a tip-affinity column (PhyTip Cat# PTR-92-20-01). HPIP was performed. Input consisted of platelet cytosol obtained from AD patients. Input and eluate samples were analyzed by 4-20% gradient SDS/PAGE followed by silver staining. TDP-43 was pull down and no IgG leach was observed. **(C)** Two tip-affinity columns were prepared using non-immune rabbit IgG (left lane) and anti-TDP-43 antibody (rabbit) (right lane) were cross-linked. The tip-affinity columns were used for immunoisolating TDP-43from platelet cytosols obtained from AD patients. No TDP-43 signal was obtained from non-immune IGG column whilst anti-TDP-43 antibody column pulled down TDP-43 proteins. Non-immune IgG column served as negative control.

**Supplemental Figure-2: Disorder enhanced phosphorylation predictor (DEPP)**. DEEP predictor revealed that the most of the phosphorylation events occur between amino acids 359-410 (framed region).

**Supplemental Figure-3: Super phosphorylated TDP-43**. Approximately 230 kDa protein peak (right-pink color) in platelet cytosols obtained from ALS patients. Anti-phosphorylated (S409/410) TDP-43 antibody was used for CEI assay. We did not observed the same peak when we performed CEI assay wit pan anti-TDP-43 antibody.

**Supplemental Table-1: Programming steps for fully automated multi-channel pipette**. These programming steps were optimized for anti-TDP-43 antibody. Users be advised that each antibody has different binding kinetics onto affinity matrix; therefore, antibody of interest need to be optimized.

## Notes

### Competing Interest Statement

The authors have declared no competing interest.

### Summary of Updates

Missing table now included in Supplementary File. Logo of FRONTIERS is now removed. Frontiers advised me that I can submit this article to bioRxiv without FRONTIERS logo.

