## Supplemental figures for "The post-translational modification profile of TAR DNA-Binding Protein (TDP-43) in platelets of patients with Alzheimer’s disease: An exploratory study for blood-based biomarker development"

**Supplemental Figure-1**

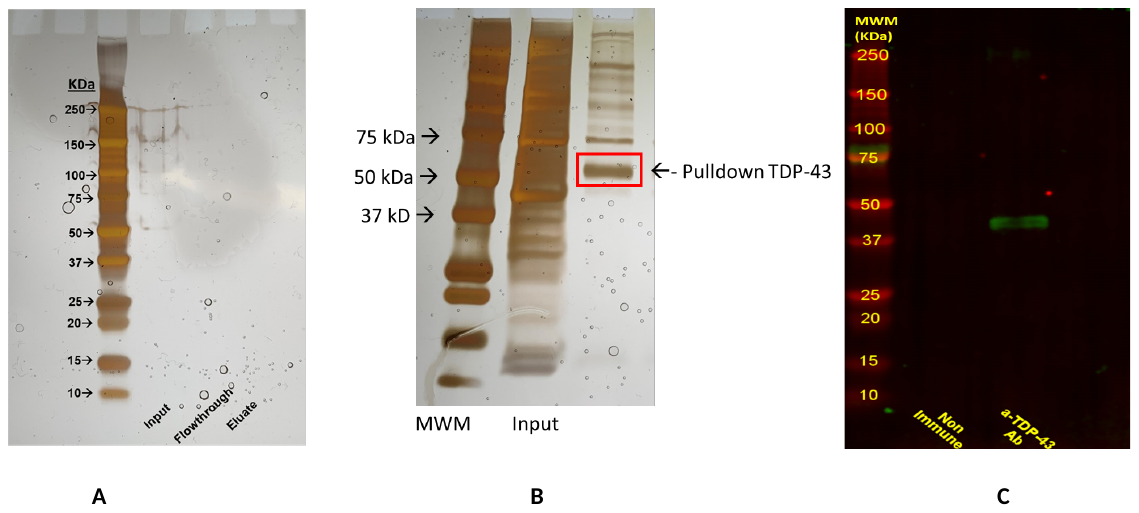

Eluate

**Supplemental Figure-2**

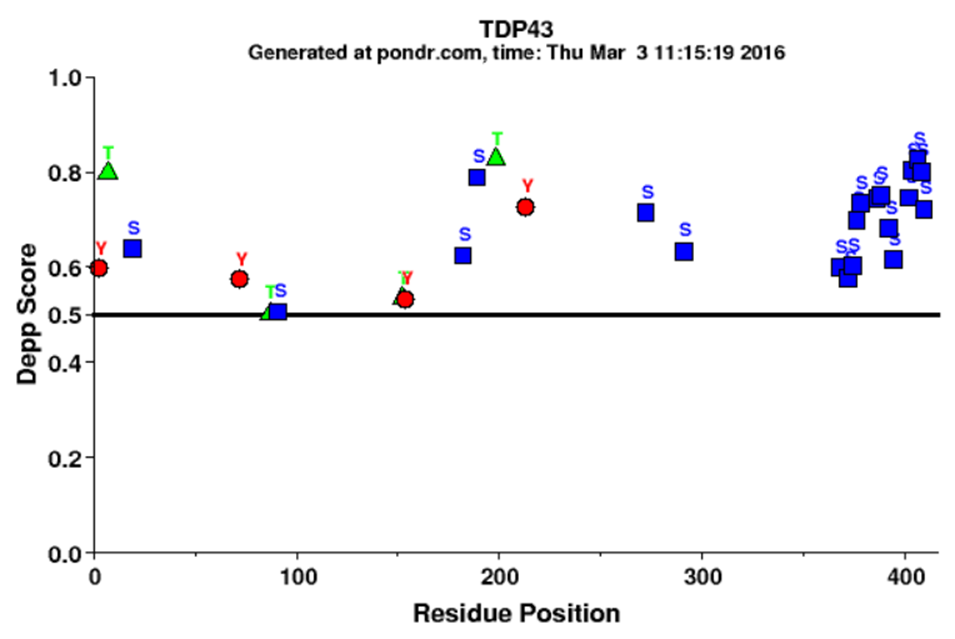

**Supplemental Figure-3**

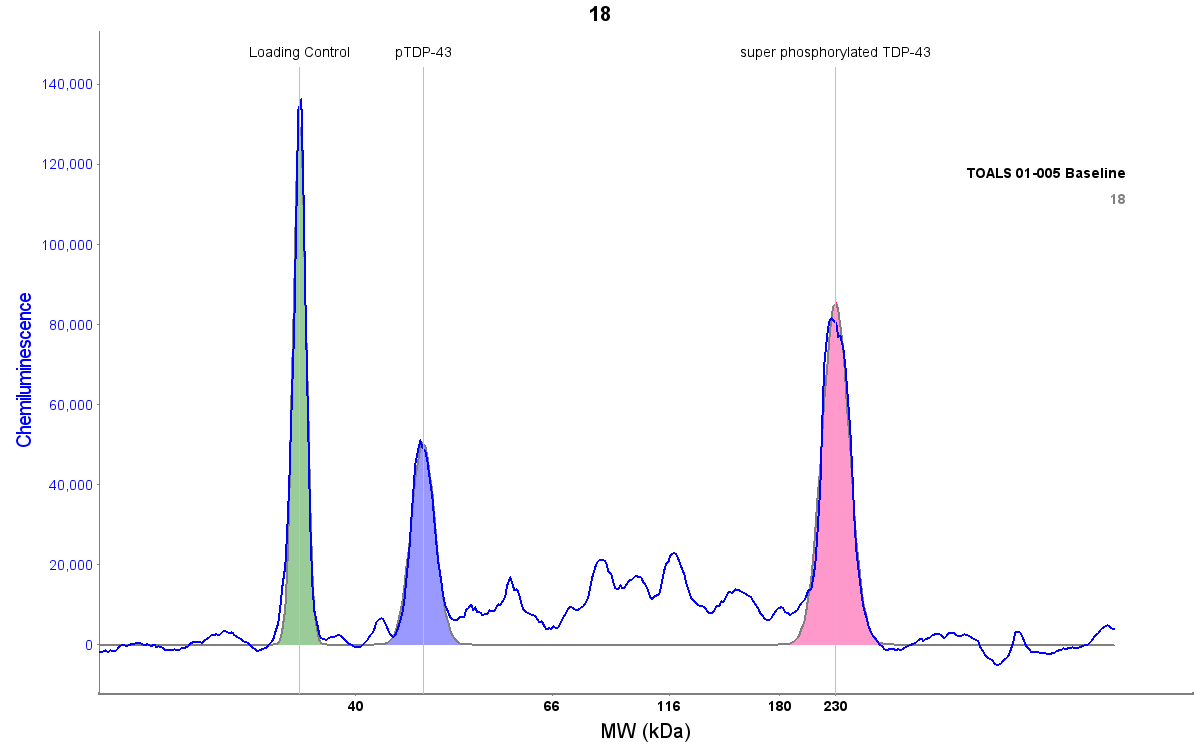

**Supplemental Table-1**

**Electronic Pipette Program for Cross-Linking Steps**

|  | Equilibrium | Capture | Wash | Eluate |
| --- | --- | --- | --- | --- |
| Volume | **100 µL** | **100 µL** | **100 µL** | **100 µL** |
| Speed | **Medium** | **Medium** | **Medium** | **Medium** |
| Aliquot | **2** | **3** | **1** | **1** |
| Cycle | **6** | **2** | **14** | **10** |

**Electronic Pipette Program for HPIP Steps**

|  | Equilibrium | Capture | Wash | Eluate |
| --- | --- | --- | --- | --- |
| Volume | **100 µL** | **100 µL** | **100 µL** | **100 µL** |
| Speed | **Medium** | **Medium** | **Medium** | **Medium** |
| Aliquot | **2** | **1** | **2** | **1** |
| Cycle | **2** | **12** | **2** | **4** |
